## Supplementary Figures for "Discovery of lipid binding sites in a ligand-gated ion channel by integrating simulations and cryo-EM"

**Zivanov J**, Nakane T, Scheres SH. A Bayesian approach to beam-induced motion correction in cryo-EM single-particle analysis. IUCrJ. 2019; 6(1):5–17.

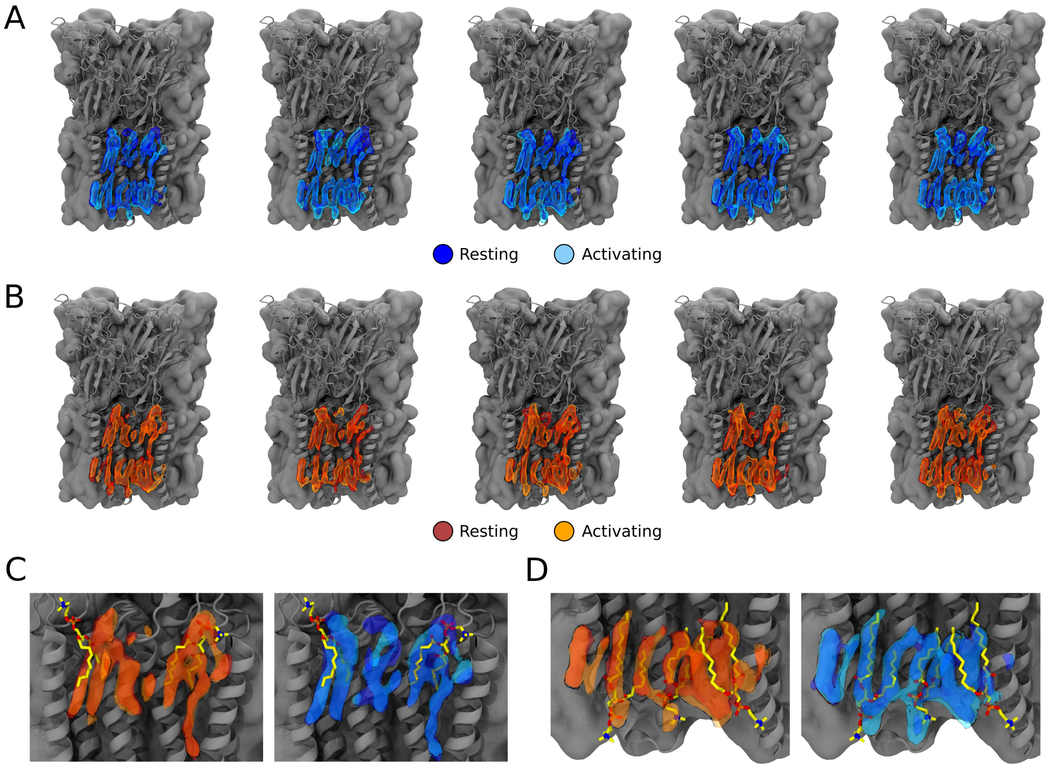


**Figure 1—figure supplement 1. Occupational lipid densities from simulations for all five subunits and overalys with built cryo-EM lipids.** (A) Lipid occupational densities obtained from simulation frames of open states at resting (dark blue) and activating (light blue) conditions for each of the five ion channel subunits. B) Lipid occupational densities obtained from simulation frames of closed states at resting (red) and activating (orange) conditions for each of the five ion channel subunits. C) Overlay of computational densities of the outer leaflet with built lipids from the cryo-EM structure (yellow sticks). D) Overlay of computational densities in the inner leaflet with built lipids from the cryo-EM structure (yellow sticks).

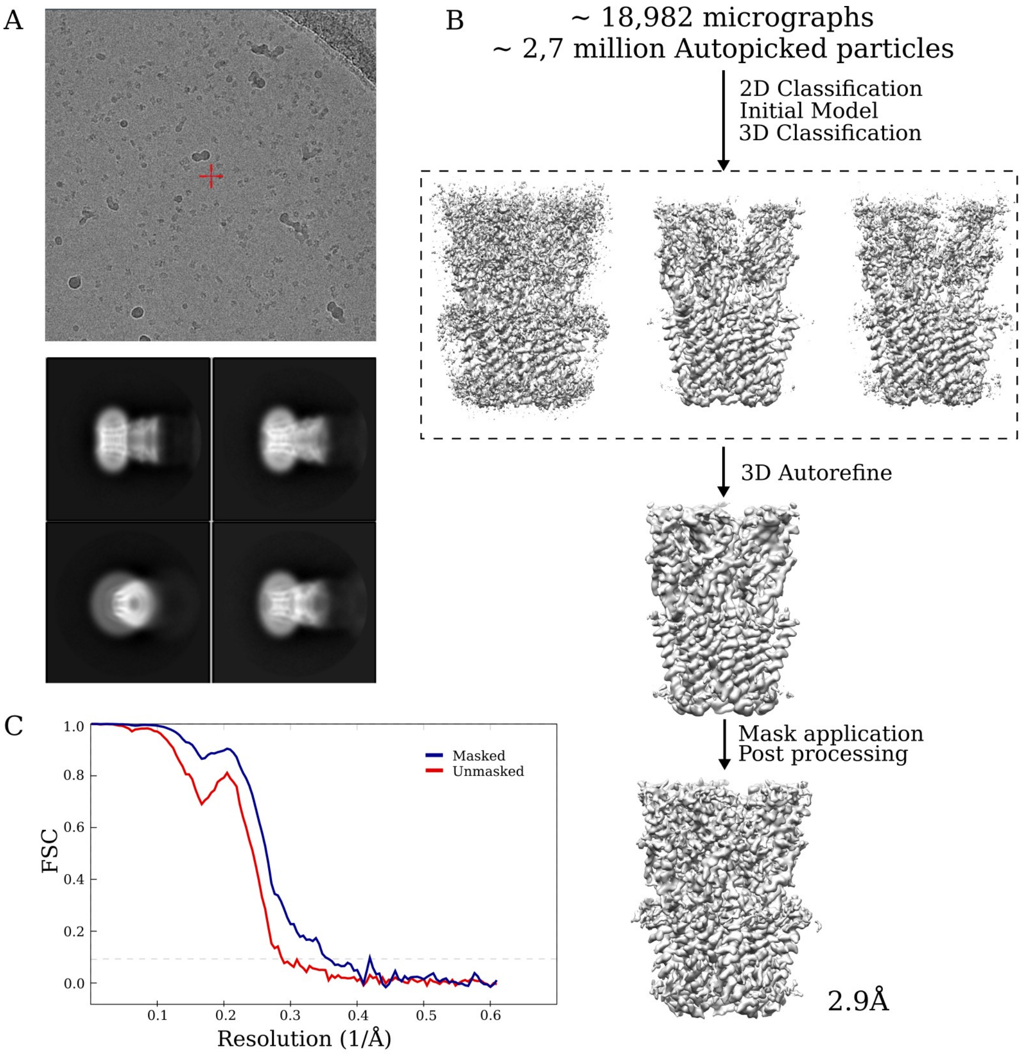


**Figure 1—figure supplement 2. Cryo-EM data and processing pipeline.** (A) Representative micrograph from a dataset collected on a Titan Krios, showing detergent-solubilized GLIC particles (*top*. Representative 2D class averages at 0.82 Å/px in a 256 x 256 pixel box and a 180 Å mask *bottom*. (B) Overview of the cryo-EM processing pipeline for merged GLIC data. (C) FSC curves for unmasked (*red*) and masked (*blue*) map.

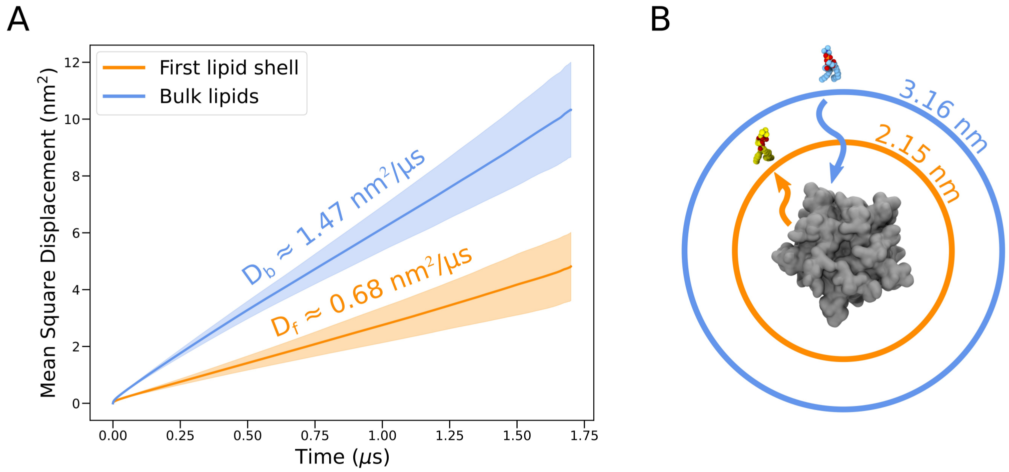


**Figure 2—figure supplement 1. Lateral diffusion of lipids initially with and without contact to the protein surface.** (A) Mean square displacements for lipids initially in the first lipid shell (<4Å from the protein surface, orange) and bulk lipids (>4Å from the protein surface, blue), averaged over all trajectories. Lighter bands represent standard deviations. Self-diffusion coefficients for lipids of the first shell, $D_f$, and bulk lipids, $D_b$, are shown above each mean square displacement line. (B) Illustration of average distances traveled by lipids initially in the first lipid shell (orange) and bulk (blue) by the end of individual simulations (1.7 μs). Distances apply in all directions of the membrane plane but are here shown as radial displacements with respect to the surface of the protein.

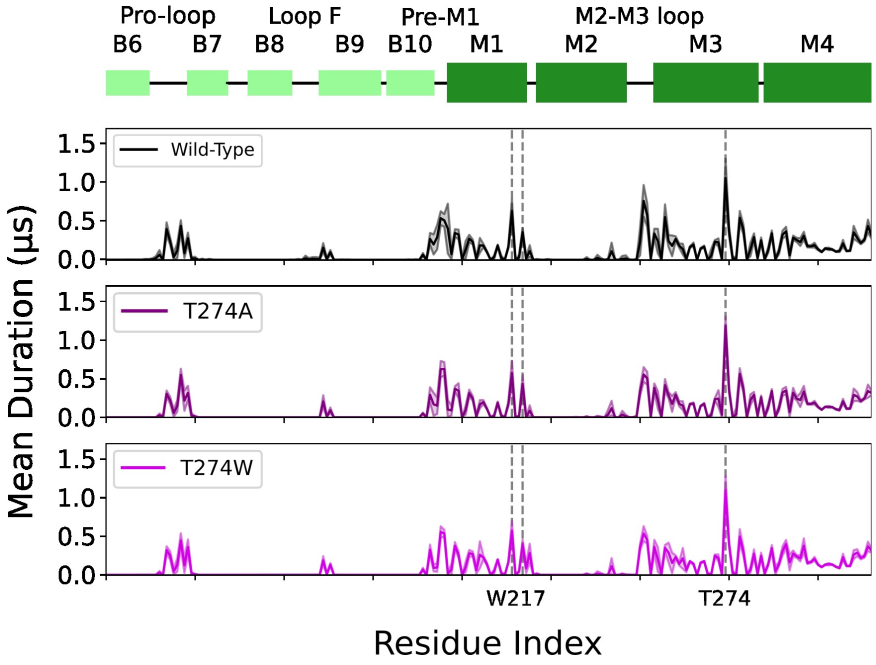


**Figure 3—figure supplement 1. W217 and T274 mutants mean duration times compared to wild type.** Mean duration time ($\mu$s) of lipid interaction time for GLIC wild type (black, *top row*), T274A mutant (purple, *middle row*) and T274W mutant (light purple, *bottom row*).

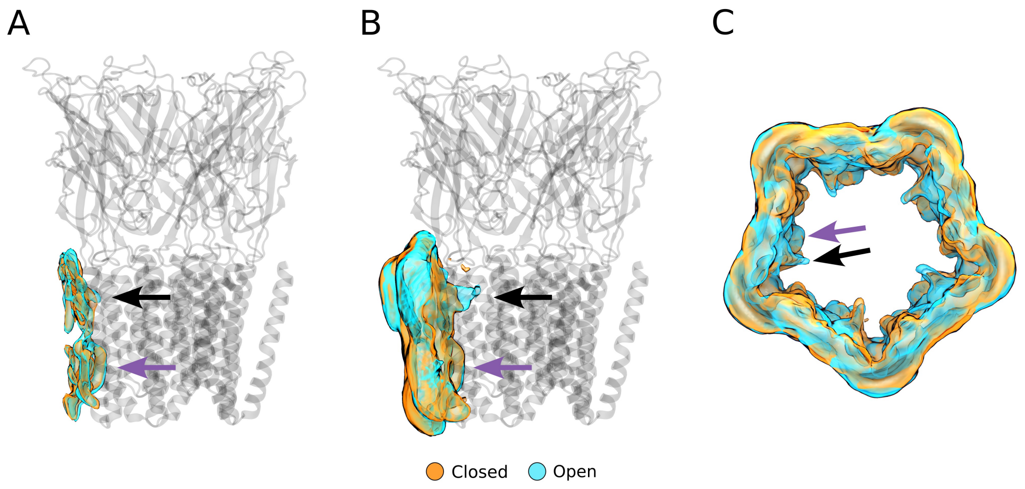


**Figure 4—figure supplement 1. Alternative views of the computationally derived lipid densities at activating conditions.** (A) Side view of a single-subunit lipid density with >40% occupancy. (B) Side view of a single-subunit lipid density with >10% occupancy. (C) Top view of the full lipid density with >10% occupancy. Black arrows indicate the outer-leaflet complementary intrasubunit lipid site, while purple arrows indicate the inner-leaflet intersubunit site.

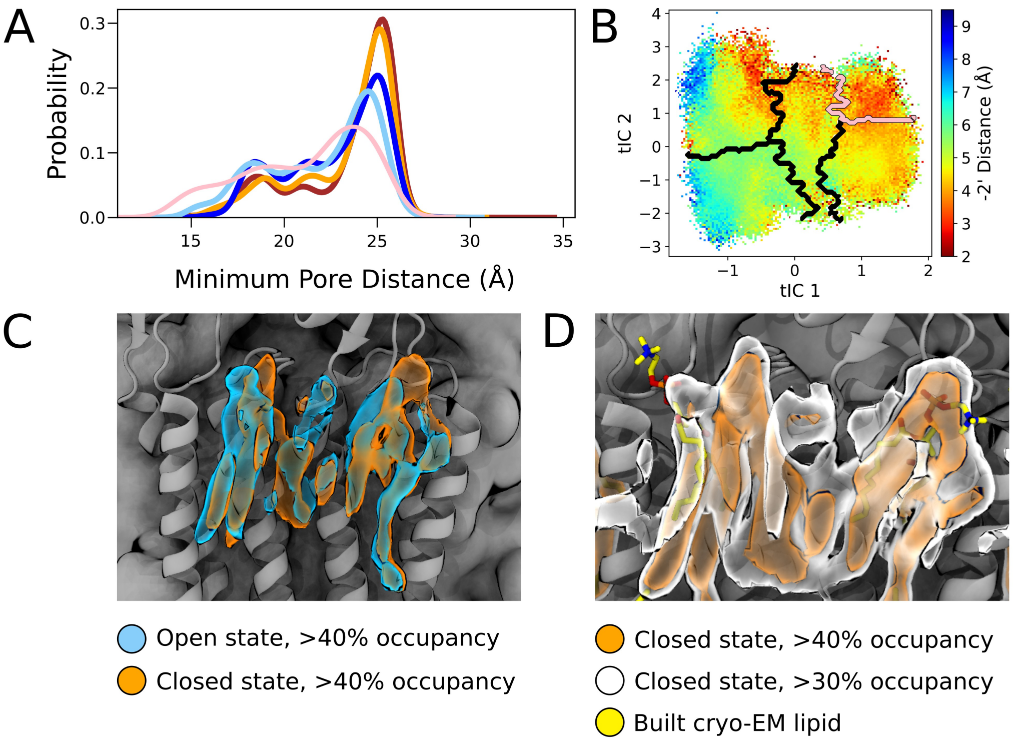


**Figure 4—figure supplement 2. Comparisons of state-dependent lipid interactions in the outer transmembrane domain.** (A) Radial distributions of nearest lipid atom from the upper pore, with a presumed pre-desensitized state displaying the highest probability of lipid tail penetration (pink), followed by open states (dark blue, light blue) and closed states (red, orange). (B) Radial distance to the -2' gate projected onto the tIC space at activating conditions. The presumed pre-desensitized state is marked in pink. (C) Lipid densities derived from simulations with >40% occupancies, representing open (blue) and closed (orange) states. (D) Lipid densities from closed state simulation frames with >40% occupancy (orange) and >30% (white). Built cryo-EM lipids are shown in yellow.

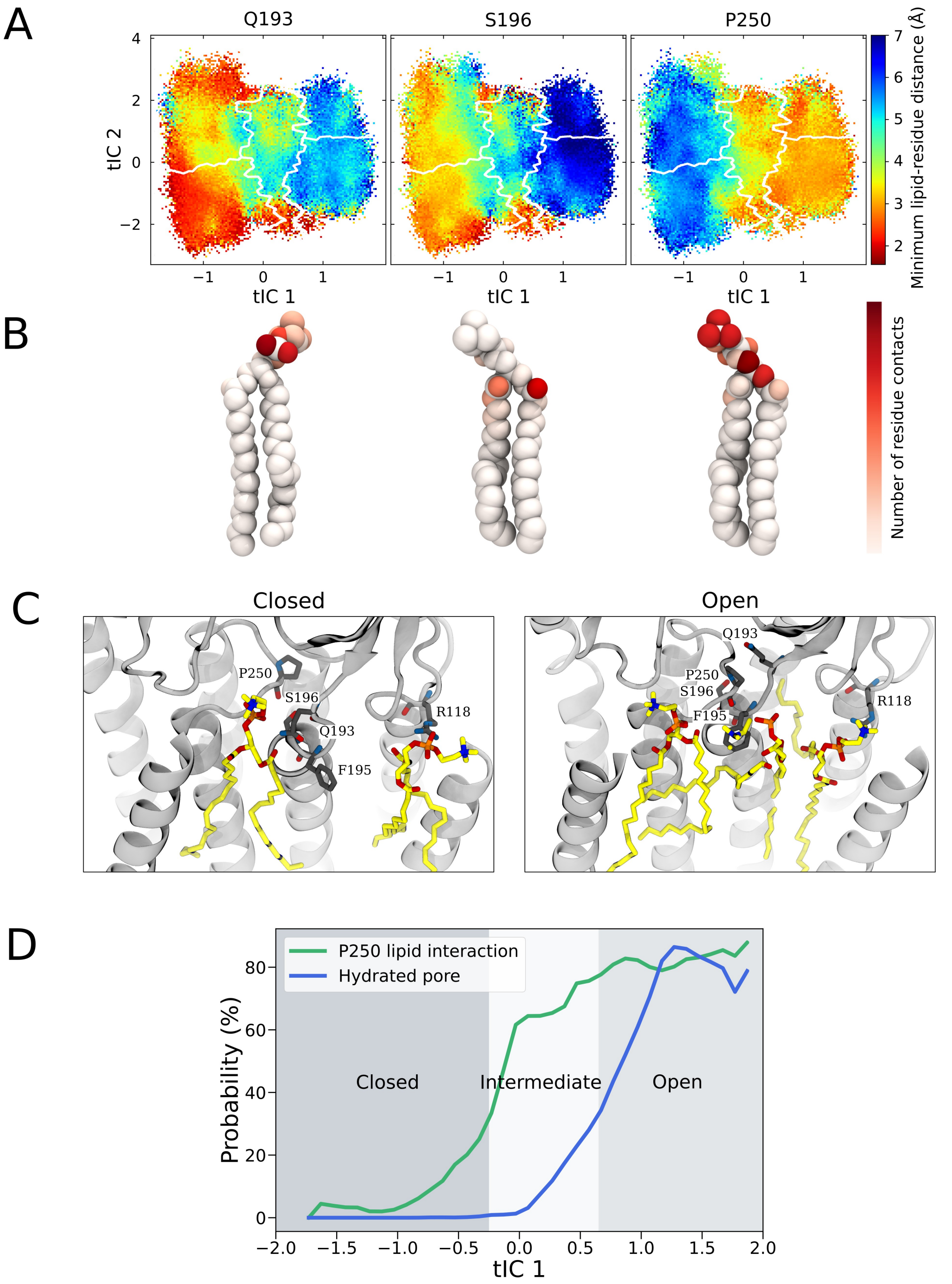


**Figure 4—figure supplement 3. State-dependent lipid interactions and conformational changes in the upper transmembrane domain.** (A) Minimum residue-lipid distance projected onto the activating condition free energy landscape for residues Q193, S196, and P250. (B) The normalized number of residue-lipid atom contacts displayed for each of the residues in (A), with red areas highlighting atoms with close interactions with the residue. (C) Representative GLIC open and closed state simulation frames with the highest correlations between lipid positions and the computational occupancies (***Figure 4***B). (D) The probabilities of P250-lipid contact formation (<4 Å) and pore hydration (>20 water molecules in the pore) projected onto the main reaction coordinate describing the gating process.

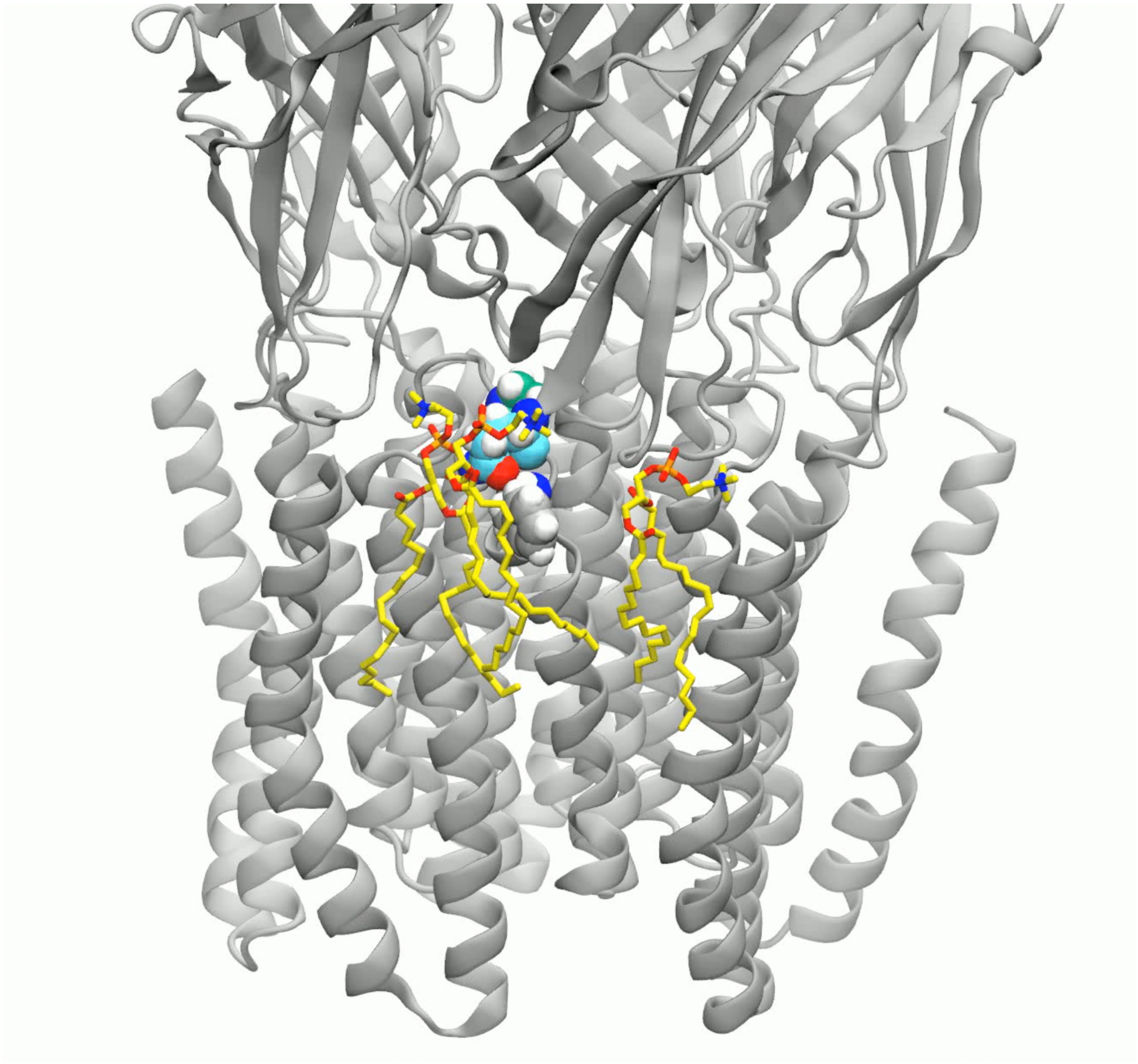

**Figure 4—Video 1. Morph between open and closed conformations showcasing changes in lipid interactions of the outer leaflet.** A linear interpolation between open and closed ion channel conformations and representative binding poses of lipids in the outer leaflet. P250 is marked by green carbon atoms, S193 by blue carbon atoms, and F195 in gray space-filling representation. Lipids are shown as yellow carbon atoms in a stick representation, and the protein as a gray cartoon representation. Oxygens are red, phosphorus orange, nitrogen blue, and hydrogens white.
